# Genetic context alters central nervous system compartment dependent responses to lipopolysaccharide

**DOI:** 10.64898/2026.07.06.736823

**Authors:** Tejas Temker, Michael MacLean, Kelly J. Keezer, Kristen D. Onos, Richard T. Libby, Gareth R. Howell

**Affiliations:** The Jackson Laboratory, Bar Harbor, ME, 04609, USA; Department of Ophthalmology, Flaum Eye Institute, University of Rochester Medical Center, Rochester, NY, USA; Neuroscience Graduate Program, University of Rochester Medical Center, Rochester, NY, USA; The Center for Visual Sciences, University of Rochester, Rochester, NY, USA; School of Graduate Biomedical Sciences, Tufts University School of Medicine, Boston, MA, 02111, USA; Graduate School of Biomedical Sciences and Engineering, University of Maine, Orono, ME, 04469, USA

## Abstract

Systemic inflammation drives neurodegeneration, yet its differential effects across neural tissues and genetic backgrounds remain poorly understood. We performed RNA-sequencing on brain, optic nerve head (ONH), and retina from four genetically diverse mouse strains (B6, CAST, NZO, WSB) following lipopolysaccharide (LPS)-induced systemic inflammation. The ONH mounted the largest response to LPS (9510 DEGs), followed by retina (5152) and brain (4586). A conserved core of 1444 DEGs across all tissues was enriched for innate immune and acute-phase pathways. Tissue-specific responses were apparent; the retina downregulated phototransduction and visual perception genes; ONH exhibited bidirectional remodeling with upregulated proteasome and ribosome biogenesis and suppressed lipid metabolism and lysosomal function; yet the brain displayed no significant pathway level enrichment. Genetic background strongly modulated the LPS response across the three tissues; the retina exhibited the greatest strain-dependent divergence. Interestingly, differing genetic context affected the ONH response to LPS the least despite its markedly larger response to LPS overall. In totality, both genetic and physical context dictate the neuroinflammatory response to LPS.

## Introduction

Inflammation in humans is a key driver in the development and progression of diseases of the brain and the eye. Systemic inflammation can disrupt the blood-brain and blood-retina barrier promoting neurodegenerative processes. Bacterial products including lipopolysaccharide (LPS) strongly activates immune cells, including central nervous system (CNS) resident microglia, triggering a potent neuroinflammatory response [1]. The brain and retina both are susceptible to inflammatory insults including inflammation induced by peripheral LPS [2]. The optic nerve head (ONH) is a specialized glial-rich region of optic nerve thought to play a critical role in the development and progression of glaucomatous neurodegeneration [3]. The ONH is thought to be a site of early injury in glaucoma [4]. Thus, it is important to understand how these distinct CNS compartments respond to systemic inflammation to identify shared and compartment-specific neurodegenerative mechanisms for increasingly common eye and brain diseases such as diabetic retinopathy, glaucoma, and Alzheimer’s disease (AD). Most studies have examined individual CNS-regions in isolation, making it difficult to compare inflammatory responses to identify shared and unique processes.

There is substantial influence of genetics on the development of eye and brain diseases. In certain cases, some single nucleotide polymorphisms have been associated with differential risk patterns for eye and brain diseases including glaucoma and AD [5, 6]. While not entirely clear what underlies this difference in risk, there is some indication it could be driven by differences in inflammatory processes [5]. The collaborative cross (CC) and diversity outbred (DO) mouse populations were generated to enable testing and mapping of genetic context on distinct phenotypes and expression patterns [7, 8]. We recently provided evidence that the founders of the CC and DO mouse populations exhibit differences in retinal aging phenotypes at transcriptional and protein levels [9]. This work strongly suggested genetic context can dictate CNS molecular aging phenotypes. In fact, we previously reported that New Zealand Obese (NZO) and Watkins Star Line B (WSB) mice of exhibit age-dependent loss of retinal ganglion cells (RGCs) and photoreceptors respectively [9]. However, it was not clear if these genetic-context driven differences in retinal aging intersect with inflammatory processes nor if this extends beyond the retina to the ONH or brain.

To address the question of how genetic context and location affects inflammation, we profiled brains, ONH, and retina samples from young 4-month-old C57BL/6J (B6), NZO, WSB, and CAST/EiJ (CAST) mice treated systemically with LPS or PBS vehicle control. This age was chosen as it is prior to RGC or photoreceptor loss in either NZO or WSB mice [9]. We identified that compartment- and genetic context-significantly impacted the response to systemic LPS. There was a shared response across all three CNS compartments to LPS exemplified by genes associated with innate and anti-viral immune responses. However, as expected, there were also tissue specific changes such as decreased gene expression profiles associated with photoreceptors in response to LPS in the retina. Further, genetic context modified the LPS response. Altogether, we have generated a high-depth dataset, available through a webtool that is broadly relevant to neuroinflammatory brain and eye diseases. Our inclusion of multiple strains provides additional genetic variability that may reflect, in part, human neuroinflammatory responses.

## Methods

### Animals and experimental design

All animal experiments were approved by the Institutional Animal Care and Use Committee at The Jackson Laboratory. Four-month-old female mice from four genetically diverse inbred strains were studied: C57BL/6J (B6; JAX 000664), CAST/EiJ (CAST; JAX 000928), NZO/HlLtJ (NZO; JAX 002105) and WSB/EiJ (WSB; JAX 001145). Lipopolysaccharide from *Escherichia coli* O111:B4 (Sigma-Aldrich, L3012) was dissolved in sterile PBS at 2.5 mg/ml. Mice received a single intraperitoneal injection of LPS (5 mg/kg) or an equivalent volume of sterile PBS. At 16h post-injection, mice were anesthetized with ketamine and xylazine and perfused transcardially with PBS. Brains were removed, hemisected and one hemibrain was snap frozen. Eyes were enucleated and then ONH and retinas were dissected and all tissues were stored at −80 °C until processing.

### RNA isolation

Dissected hemibrains were homogenized in MR1 buffer (Macherey-Nagel, 744351.125) using a gentleMACS dissociator (Miltenyi Biotec), while retina and ONH were lysed and homogenized in TRIzol Reagent (ThermoFisher). Total RNA was isolated from using the NucleoMag RNA kit (Macherey-Nagel, 744350.4) on a KingFisher Flex purification system (Thermo Fisher Scientific), according to the manufacturer’s instructions. RNA quantity and quality were evaluated using a NanoDrop 8000 spectrophotometer (Thermo Scientific) and Agilent RNA 6000 Nano and Pico assays. Only samples with an RNA integrity number greater than 7 were used for library preparation.

### Library preparation and RNA sequencing

Strand-specific RNA-seq libraries were generated using the KAPA RNA HyperPrep Kit with RiboErase (Roche Sequencing and Life Science, 08098131702) according to the manufacturer’s protocol. Library quality and concentration were assessed using Agilent D5000 ScreenTape and the Qubit dsDNA HS assay (Thermo Fisher Scientific, Q32851), respectively. Samples were sequenced on an Illumina NovaSeq 6000 using the S4 Reagent Kit v1.5 (Illumina, 20028313) at the Genome Technologies Core at The Jackson Laboratory, yielding approximately 60 million 150-bp paired-end reads per sample.

### RNA-seq data processing and alignment

Raw FASTQ files were processed using standard quality-control procedures. High-quality read pairs were aligned with STAR v2.7.9a [10] to a custom pseudogenome generated by incorporating strain-specific single-nucleotide variants and short insertions and deletions from Mouse Genomes Project REL-1505 [11] into the mouse reference genome (GRCm38/mm10) using g2gtools version 0.2.7 (https://github.com/churchill-lab/g2gtools). Custom genomes were generated for NZO/HlLtJ, CAST/EiJ and WSB/EiJ mice, whereas reads from C57BL/6J mice were aligned to the unmodified mm10 reference. Read counts were quantified with the featureCounts function in Subread v1.6.3 [12, 13] using Ensembl Release 68 transcriptome annotation as the reference [14].

### Differential expression analysis

Differential expression analyses were performed using edgeR [15] within the R environment via the quasi-likelihood F-test framework (glmQLFTest) with robust dispersion estimation. Library sizes were normalized using the trimmed mean of M-values (TMM) method.

Two complementary analyses were performed. First, brain, retina, and ONH were analyzed separately to estimate the tissue-specific LPS response as the unweighted average LPS-versus-PBS effect across B6, CAST, NZO, and WSB. Second, strain-dependent responses relative to B6 were tested within each tissue using interaction contrasts of the form (Strain X LPS - Strain X PBS) - (B6 LPS - B6 PBS) for CAST, NZO, and WSB. These contrasts tested for differences in the LPS response relative to B6, not uniqueness across all strains. Genes were considered differentially expressed genes with adjusted *p*-value of <0.05 and log_2_FC of >0.5.

Over-representation analysis of differentially expressed genes was performed using the clusterProfiler package in R [16], testing for enrichment of Gene Ontology Biological Process [17] with all detected genes used as the statistical background.

To examine the strain-specific vulnerabilities Photoreceptor marker genes were visualized for WSB mice and RGC marker genes for NZO mice. Curated cell type-specific marker gene lists derived from Marola et al. [9] which were visualized as heatmaps. The log-CPM were normalized using the TMM and scaled across all replicates (row z-score). Similarly, to visualize the expression of microglial activation signatures, curated interferon response module (IRM) and disease-associated microglia (DAM) gene sets were examined across all three CNS regions. The log-CPM values were computed per tissue using TMM-normalized DGEList objects via edgeR, and row z-scores were calculated per gene across all replicates within each tissue.

## Results

### Global Transcriptomic Response to LPS

We used a single dose of lipopolysaccharide (LPS) to model acute systemic inflammation in four inbred mouse strains C57BL/6J (B6), CAST/EiJ (CAST), NZO/HILtJ (NZO), and WSB/EiJ (WSB). We then collected brain, optic nerve head (ONH), and retinal tissue and performed whole tissue RNA-sequencing. We performed differential expression analysis comparing LPS and PBS treated animals across all three CNS tissues to identify shared and unique mechanisms of inflammation across regions. The inclusion of four genetically distinct mouse strains better enables us to mirror, at least in part, genetic diversity present in human populations.

Multidimensional scaling (MDS) of all samples showed that the primary axis of transcriptomic variation is dictated by tissue identity, with brain, ONH, and retina forming three well-defined clusters (Supplemental Figure 1A). Within each tissue cluster, LPS and PBS treated samples appear largely intermingled along both MDS1 and MDS2 (Supplemental Figure 1B). Similarly, coloring by mouse strain shows that all four inbred strains are distributed across each tissue cluster (Supplemental Figure 1C).

LPS induced widespread transcriptomic changes across all three tissues (Figure 1A). The ONH exhibited the largest response with 9510 differentially expressed genes (DEGs). The retina showed an intermediate response with 5152 DEGs. Whereas the brain displayed the smallest, though still substantial, response with 4586 DEGs.

**Figure 1.**
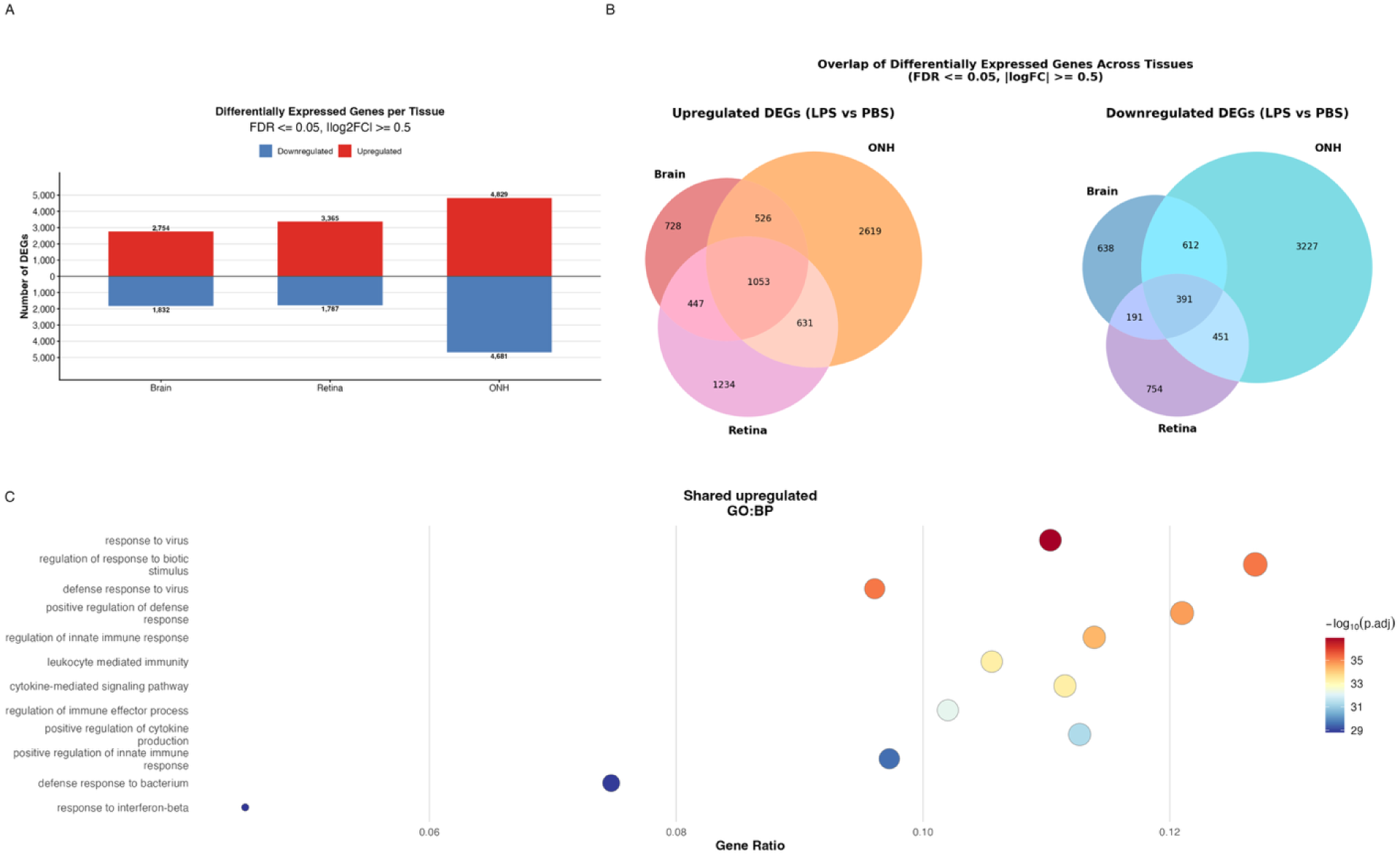
LPS induces widespread but region-divergent transcriptomic changes across CNS regions. (A) Bidirectional bar chart depicting the number of upregulated (red, above zero) and downregulated (blue, below zero) differentially expressed genes (DEGs) in each tissue following systemic LPS challenge (brain: LPS n=14, PBS n=11; ONH: LPS n=18, PBS n=13; retina: LPS n=19, PBS n=17). (B) Three-way Venn diagrams illustrating the overlap of DEGs (FDR ≤ 0.05, |logFC| ≥ 0.5) across brain (LPS n=14, PBS n=11), ONH (LPS n=18, PBS n=13), and retina (LPS n=19, PBS n=17), shown separately for upregulated (left) and downregulated (right) genes (LPS vs. PBS).

To determine the extent to which the LPS response is shared or tissue-specific, we compared DEGs across tissues (Figure 1B). A set of 1444 DEGs were detected in all three tissues, suggestive of a conserved LPS-induced gene signature. The shared up-regulated genes were enriched for innate immunity and viral response GO biological processes (Figure 1C). The shared down-regulated genes were not enriched for any GO biological process, suggesting that LPS reduced gene expression across diverse processes. Interestingly, 1082 genes were shared between the ONH and retina, 1138 between the brain and ONH, and 638 between the brain and retina. The ONH contained the largest set of tissue-specific DEGs (5846 genes), more than three times the retina (1988) and brain (1366).

The shared core response was dominated by canonical innate immune and acute-phase mediators. However, the most strongly upregulated genes differed by tissue: *Saa3, Lcn2, Fpr1, Fpr2,* and *Timp1* were prominent in the ONH; *Txnip* was most strongly induced in the brain; and interferon-stimulated genes *Irf7* and *H2-T23* reached significance at lower fold-changes in the retina. (Figure 2). These findings highlight that neuroinflammatory responses to LPS are not uniform but are shaped by tissue context. To further characterize these differences, we next examined tissue-specific transcriptional programs in greater detail.

**Figure 2.**
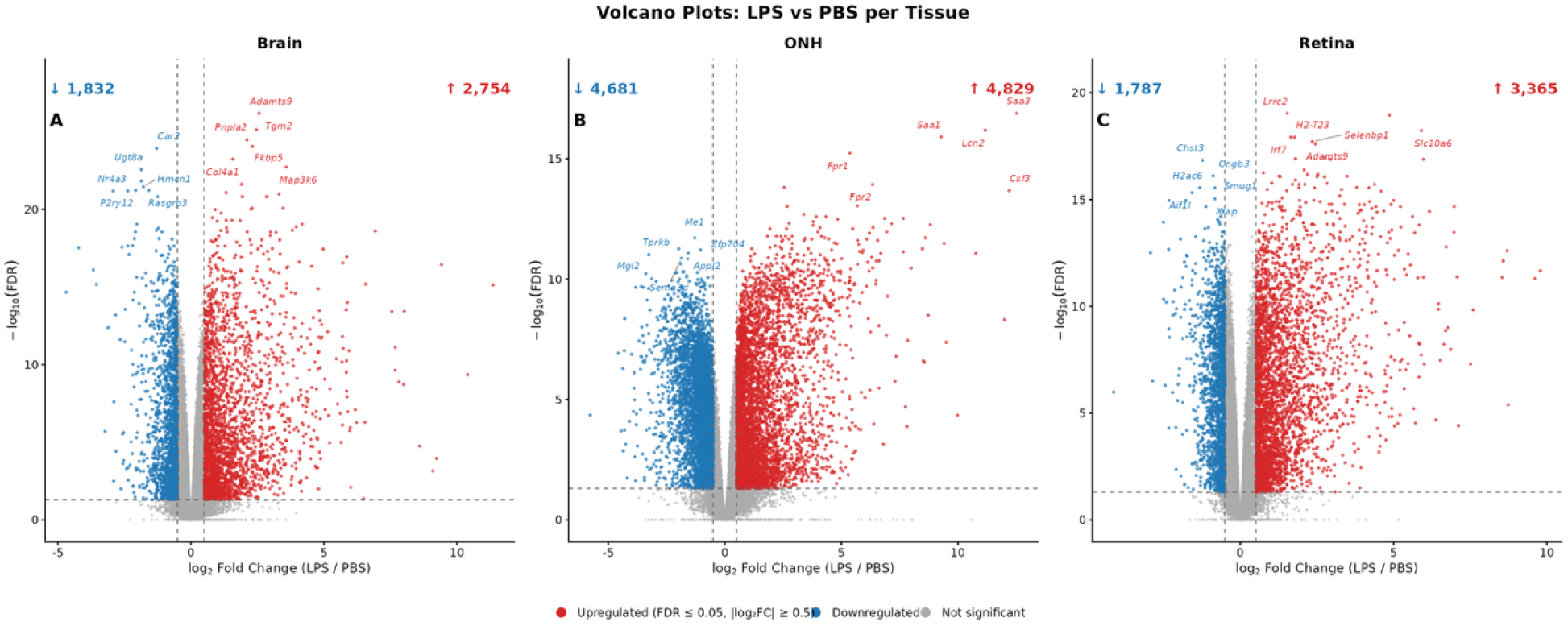
Volcano plots reveal tissue-specific patterns of differential gene expression following LPS challenge. Volcano plots of differential expression results **A.** brain (LPS n=14, PBS n=11), **B.** ONH (LPS n=18, PBS n=13), and **C.** retina (LPS n=19, PBS n=17). The x-axis represents log fold change, and the y-axis represents −log (FDR). Red points indicate upregulated genes (FDR ≤ 0.05, logFC ≥ 0.5); blue points indicate downregulated genes (FDR ≤ 0.05, logFC ≤ −0.5); grey points are not significant.

### Systemic LPS Elicits Broad but Tissue-Divergent Transcriptomic Responses

Brain-specific upregulated genes (n = 728) were enriched for GO biological process terms including visual perception and sensory perception of light stimulus. There was no significant GO biological process enrichment for the down-regulated genes.

In contrast, the retina exhibits a combination of mixed inflammation regulatory signals suggesting both activation and suppression. Retina-unique upregulated genes (n = 1234) were enriched for detection of chemical stimulus, sulfur compound transport, ossification, and biomineral tissue development (Figure 3). Retina-unique downregulated genes (n = 754) were enriched for detection of light stimulus, detection of abiotic stimulus, detection of external stimulus, visual perception, sensory perception of light stimulus, and phototransduction (Figure 4). This coordinated decrease in expression across phototransduction-related genes suggests that endotoxemia may alter retinal function consistent with earlier studies showing that systemic LPS enhances retinal degeneration in dystrophic models [18].

**Figure 3.**
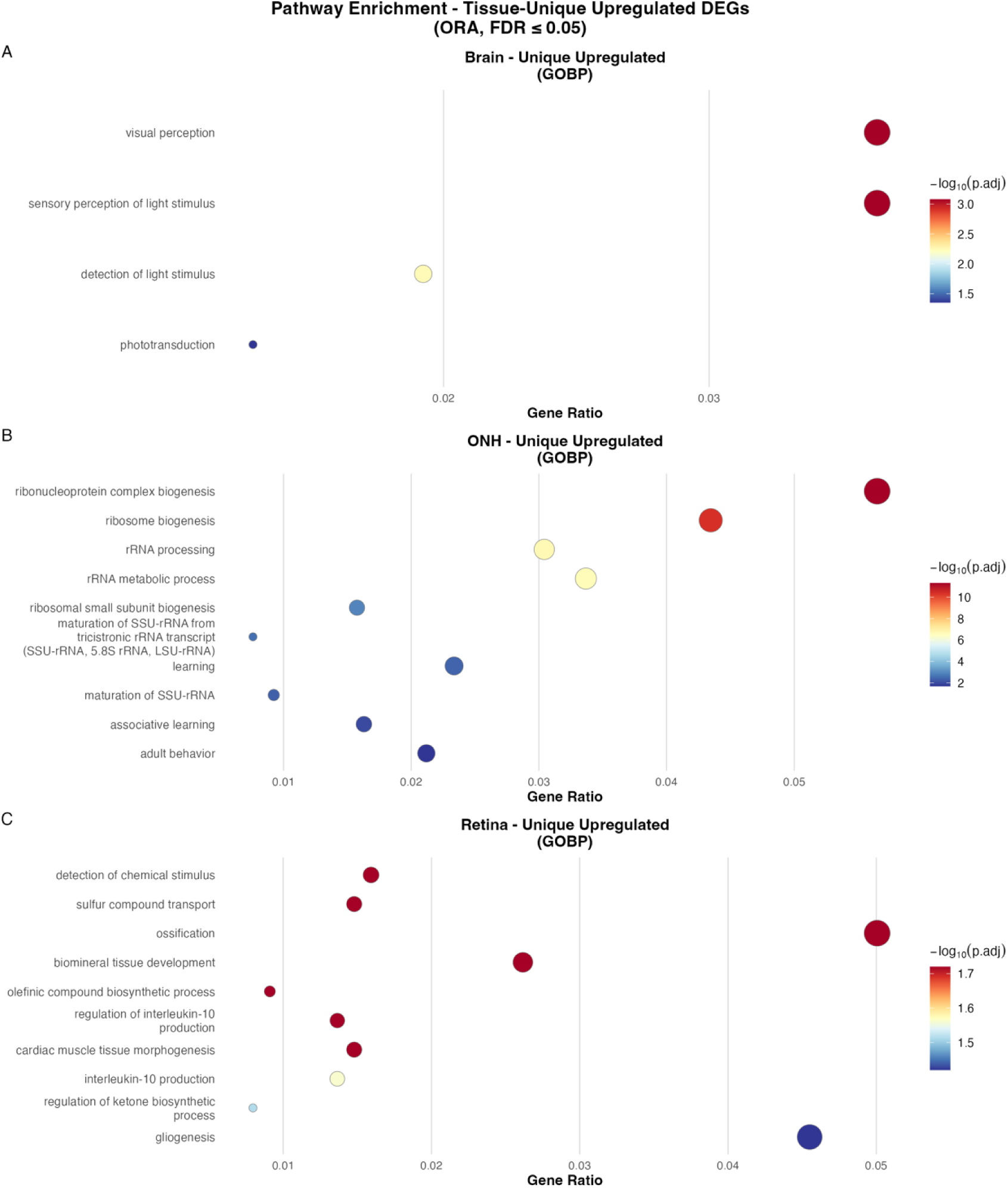
Over-representation analysis of tissue-unique upregulated DEGs reveals divergent pathway activation across CNS tissues. Dot plots depicting GO Biological Process (GOBP) over-representation analysis (ORA) results for tissue-unique upregulated DEGs in brain (top row; LPS n=14, PBS n=11), ONH (middle row; LPS n=18, PBS n=13), and retina (bottom row; LPS n=19, PBS n=17). Dot size represents gene ratio; color represents −log (adjusted P-value).

**Figure 4.**
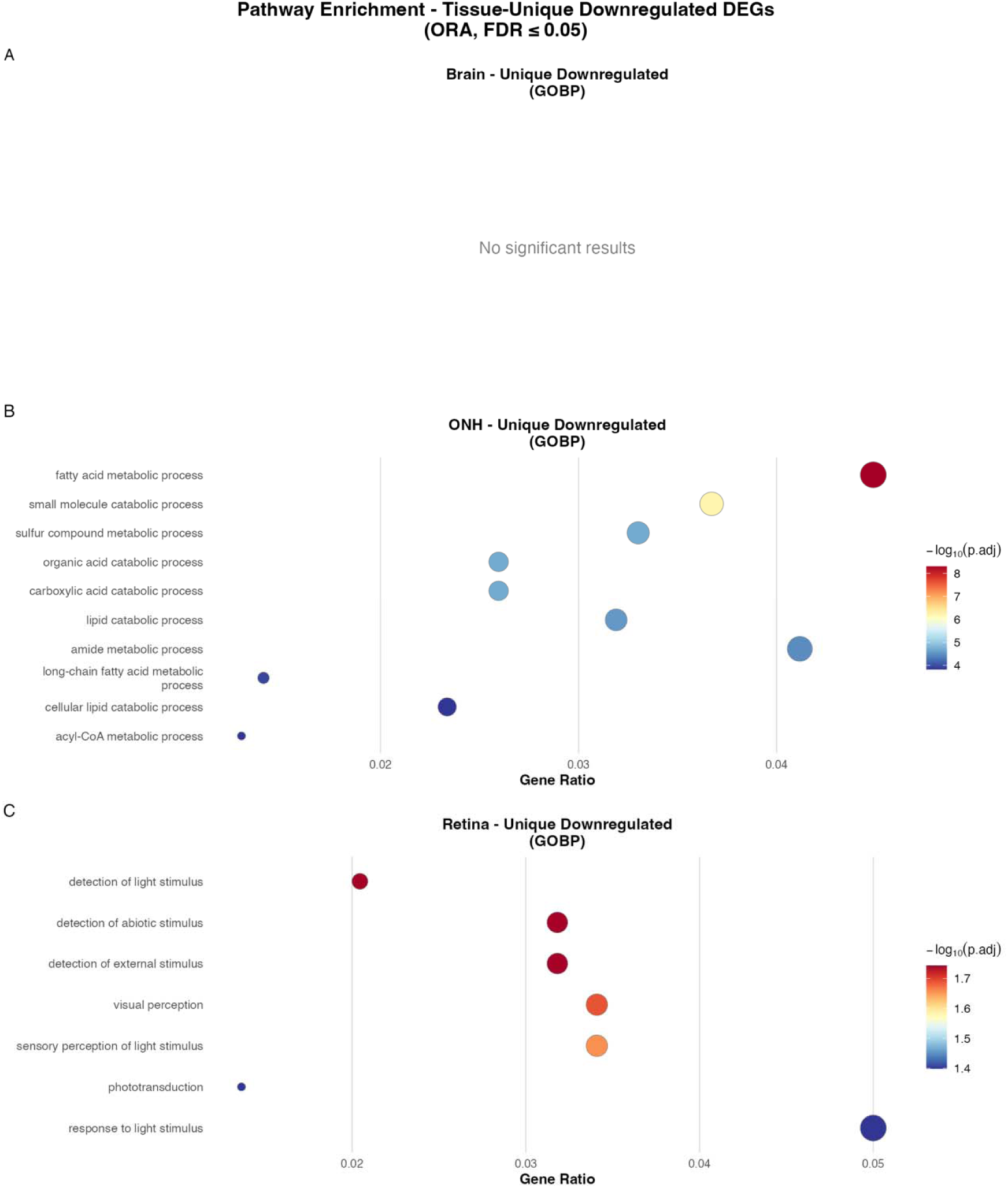
Over-representation analysis of tissue-unique downregulated DEGs reveals suppression of neuroactive signaling, phototransduction, and lipid metabolism. Dot plots depicting GOBP ORA results for tissue-unique downregulated DEGs in brain (top row; LPS n=14, PBS n=11), ONH (middle row; LPS n=18, PBS n=13), and retina (bottom row; LPS n=19, PBS n=17). Dot size represents gene ratio; dot color represents −log (adjusted P-value).

The ONH mounted the largest transcriptomic response of 9510 DEGs with a near-equal split between upregulated (4829) and downregulated (4681) genes (Figure 1A). This extensive response could be explained by features of the ONH including how its cellular composition is dominated by astrocytes, which may confer a heightened transcriptional sensitivity to systemic inflammation. Pathway enrichment of ONH-unique upregulated genes showed proteasome, ribosome biogenesis in eukaryotes, and Spinocerebellar ataxia (Figure 3). Additionally, ONH-unique downregulated genes (n = 3227) were enriched for fatty acid metabolic process, lipid catabolic process, lysosome biogenesis, pyruvate metabolism, metabolism of xenobiotics by cytochrome P450, Drug metabolism – cytochrome P450, and Regulation of lipolysis in adipocytes (Figure 4). Altogether, the ONH exhibited proteostasis and metabolic gene expression shifts in response to LPS.

Together, these tissue-specific signatures indicate that while LPS induced dysregulation of neural, metabolic, and immune gene programs across all CNS compartments, the specific functional pathways affected differed between tissues. Having described the shared and tissue-specific responses to LPS, we next asked whether these responses are further modulated by genetic context.

### Genetic context modulates the transcriptional response to LPS across CNS tissues

We have previously reported that retinal aging molecular signatures are modified by genetic context in these four mouse strains [9]. In the retina, NZO and WSB mice exhibit age-related loss in retinal ganglion cells and photoreceptors respectively [9]. Thus, to determine the impact of genetic context on the neuroinflammatory response to LPS, we compared LPS-induced gene expression changes in CAST, WSB, and NZO mice relative to B6 within each CNS region. These three strains were selected to span a range of genetic divergence from B6: CAST is a wild-derived Mus musculus castaneus strain and one of the most genetically distant Collaborative Cross founders from B6, capturing much of the known genetic variation among laboratory mice [19]; WSB is wild-derived but M. m. domesticus, representing an intermediate level of divergence; and NZO is a classical inbred strain, more closely related to B6 [19]. This design allowed us to assess whether the magnitude of the neuroinflammatory response scales with genetic distance from B6. The level of the strain-differential transcriptional response varied substantially by both strain and region (Figure 6-9).

Within the brain WSB (606 DEGs), NZO (519 DEGs), and CAST (376 DEGs) all exhibited strain-differential responses to LPS (Figure 5A). Yet, there were no significant pathway enrichment for genes associated with strain-associated LPS DEGs for WSB or CAST relative to B6 mice. NZO mice showed a modest but significant enrichment of genes showing a lower degree of LPS-induced changes in collagen biosynthetic and metabolic pathways relative to B6 mice including *Rgcc, Itga2, Ccn2, Il6ra, Arg1, Errfi1, Dicer1*, and *Tgfbr3* (Figure 5B). The overall paucity of pathway-level enrichment in the brain, despite hundreds of strain-differential DEGs, could hint that strain-differential effects are distributed across many biological processes.

**Figure 5.**
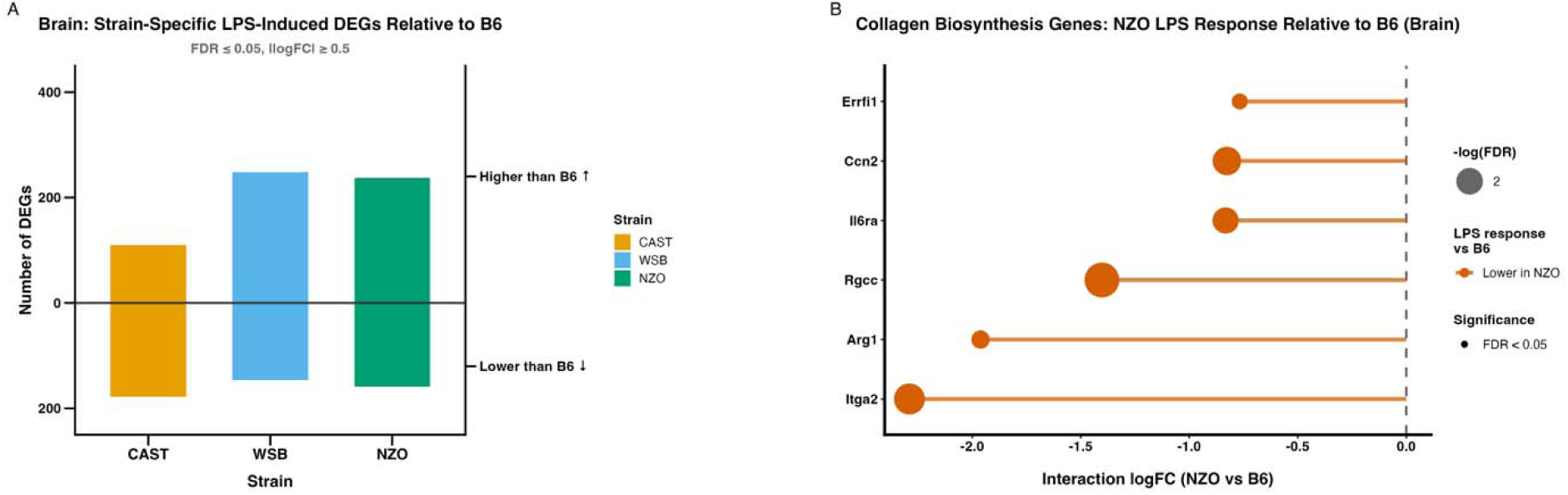
Evaluation of strain-divergent responses to LPS within the brain. (A) Bidirectional bar chart showing the number of DEGs (FDR ≤ 0.05, |logFC| ≥ 0.5) per strain relative to B6 in the brain (B6: LPS n=3, PBS n=3; CAST: n=4/2; NZO: n=3/3; WSB: n=4/3). Bars above zero indicate genes with greater LPS-induced expression relative to B6; bars below zero indicate genes with lower LPS-induced expression relative to B6. Expression of collagen biosynthesis pathway genes in brain tissue from B6 and NZO mice (n=3 per group).

In contrast, the retina exhibited large strain-divergent responses with WSB and NZO mice exhibiting 2115 and 1490 DEGs, respectively, while CAST mice exhibited 639 DEGs relative to B6 (Figure 6A). In the retina, WSB mice exhibit increased changes in the expression of genes enriched for DNA replication, mitotic spindle organization, glycosylation, and positive regulation of DNA metabolic processes relative to B6 after LPS (Figure 6B). This is alongside a modest reduction in LPS-induced changes in genes associated with endothelial cell migration. Genes more affected by LPS in NZO mice were enriched for ribonucleoprotein complex biogenesis, rRNA metabolic processes, rRNA processing, ribosome biogenesis, DNA replication and repair, photoreceptor cell development, and macroautophagy pathways (Figure 6B). DEGs associated with CAST divergent responses to LPS showed no significant enrichment in the retina despite 639 DEGs. To determine if NZO and WSB mice treated with LPS exhibit accelerated RGC or photoreceptor loss we visualized RGC and photoreceptor marker genes using a heatmap within each strain (Figure 6C-D). There were up-regulated and down-regulated genes in both NZO RGC genes and WSB photoreceptor genes in response to LPS (Figure 6C-D). Altogether, these data suggest acute systemic inflammation may modulate the neural dysfunction in these models.

**Figure 6.**
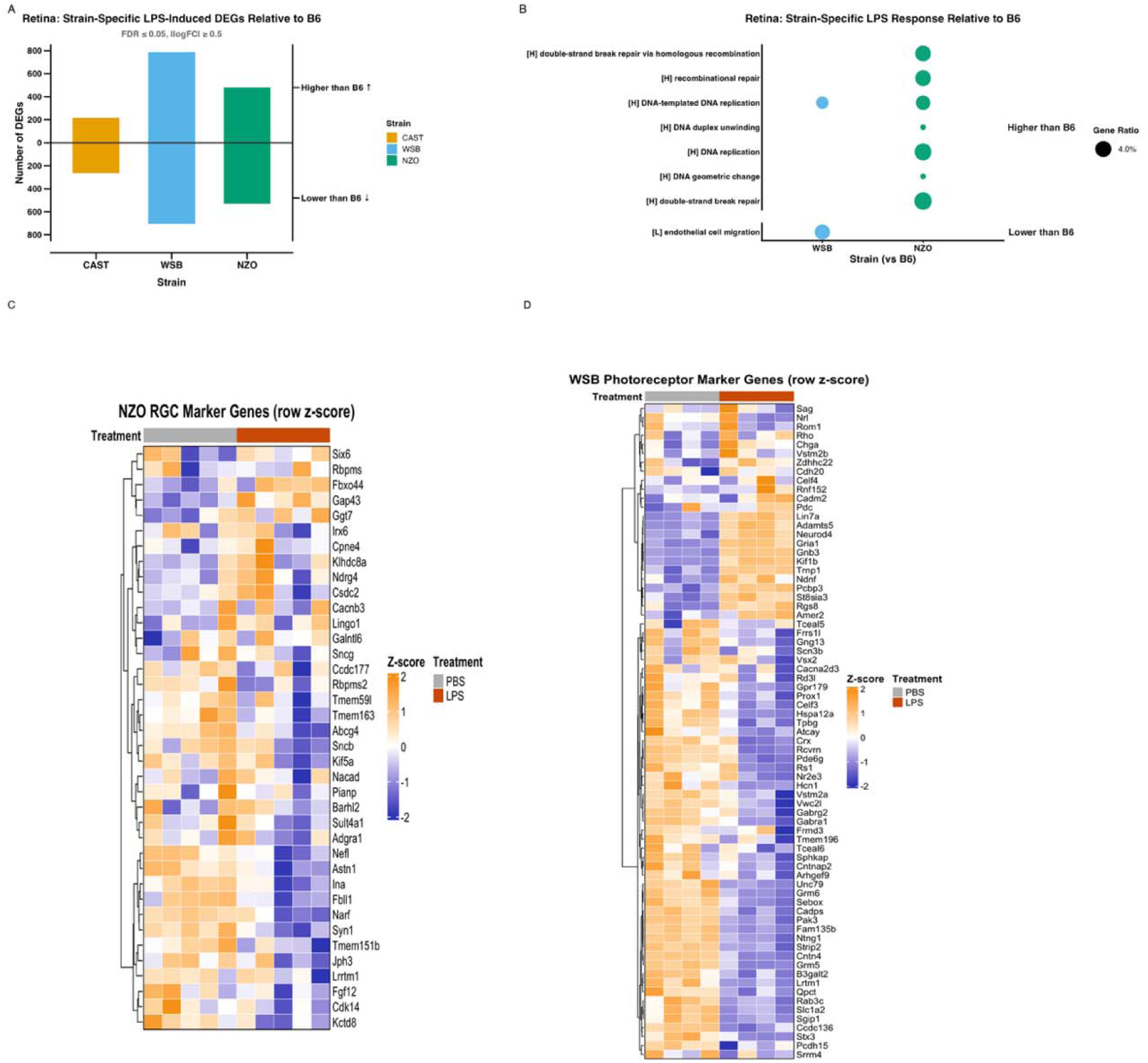
Evaluation of strain-divergent responses to LPS within the retina. (A) Dot plot depicting overrepresentation of GOBP in the retina (CAST, WSB, and NZO vs. B6) (B6: LPS n=4, PBS n=4; CAST: n=6/4; NZO: n=5/5; WSB: n=4/4). Dot size represents gene ratio (2–6%); dot color represents strains. (B) Bidirectional bar [9]chart showing the number of DEGs (FDR ≤ 0.05, |logFC| ≥ 0.5) per strain relative to B6. Bars above zero indicate genes with greater LPS-induced expression relative to B6; bars below zero indicate genes with lower LPS-induced expression relative to B6. (C) Heatmap of curated retinal ganglion cell (RGC) marker gene expression in NZO retinas (PBS n=5, LPS n=5), Expression values are log-CPM scaled across all replicates. (D) Heatmap of curated photoreceptor marker gene expression in WSB retina (PBS n=4, LPS n=4). Displayed as in (C).

The ONH showed the smallest divergent response with WSB exhibiting 415 DEGs, CAST 309 DEGs, and NZO only 106 DEGs relative to B6 (Figure 7A). Differentially responsive LPS DEGs in CAST or NZO relative to B6 mice did not exhibit any enrichment for GO Biological Pathways. However, genes more affected by LPS in WSB mice relative to B6 mice showed significant enrichment in pathways related to the endoplasmic reticulum stress responses (Figure 7B). These data suggest that WSB ONH tissue exhibits an even greater proteostatic stress response to LPS than B6 mice as this was a dominant signal in the average LPS-induced gene signature in ONH (Figure 3).

**Figure 7.**
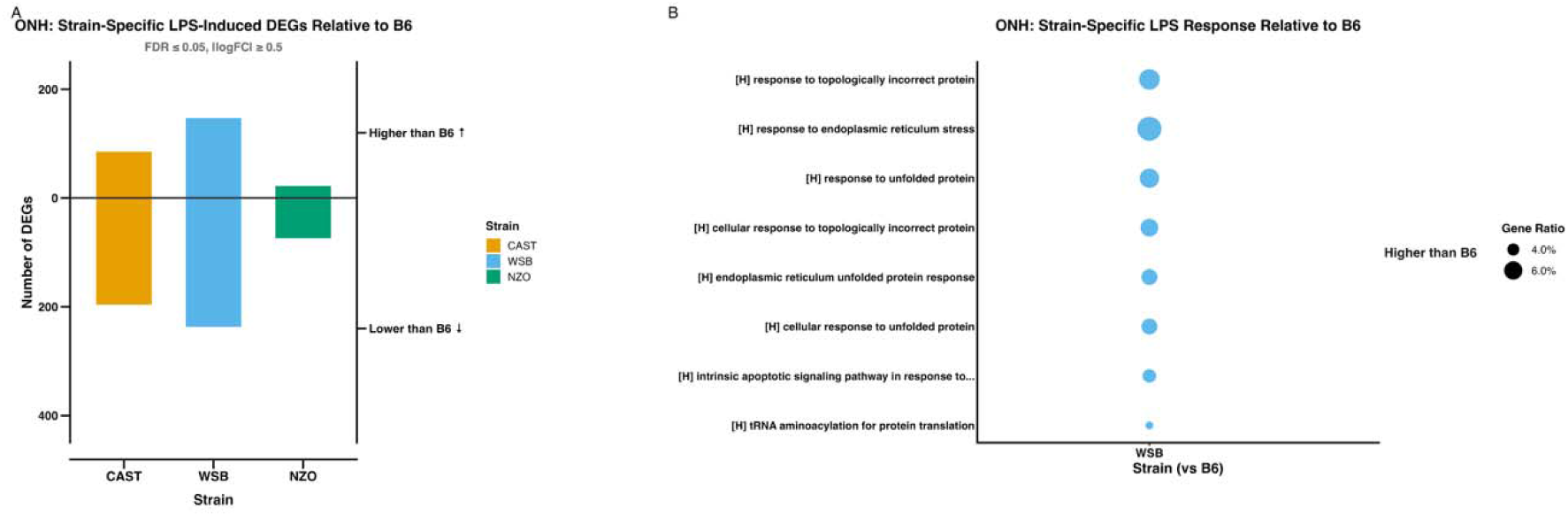
Evaluation of strain-divergent responses to LPS within the optic nerve head. (A) Dot plot depicting overrepresentation of GOBP in the ONH (B6: LPS n=4, PBS n=3; CAST: n=6/3; NZO: n=4/4; WSB: n=4/3). Dot size represents gene ratio (2–6%); dot color represents strains. (B) Bidirectional bar chart showing the number of DEGs (FDR ≤ 0.05, |logFC| ≥ 0.5) per strain relative to B6. Bars above zero indicate genes with greater LPS-induced expression relative to B6; bars below zero indicate genes with lower LPS-induced expression relative to B6.

Finally, to frame the LPS-induced response on the resident immune cells of the CNS, microglia, we examined the expression of curated interferon responding microglia (IRM) and disease-associated microglia (DAM) gene signatures across all three CNS regions and all four strains (Figure 8). DAM and IRM are two of the most common activated states in brain and eye diseases. The IRM genes were broadly and robustly induced by LPS across all tissues in all strains, consistent with a conserved interferon-driven innate immune response to systemic endotoxin. In contrast, homeostatic genes were suppressed by LPS, most prominently in the retina, suggesting a shift away from the homeostatic microglial state. DAM activation-associated genes, including *Trem2, Gpnmb, Spp1, Clec7a, Lpl,* and *Cst7* exhibited variable upregulation across regions. The retina exhibited the most pronounced LPS-driven shift in both gene sets, consistent with its greater overall transcriptional responsiveness to systemic inflammation.

**Figure 8.**
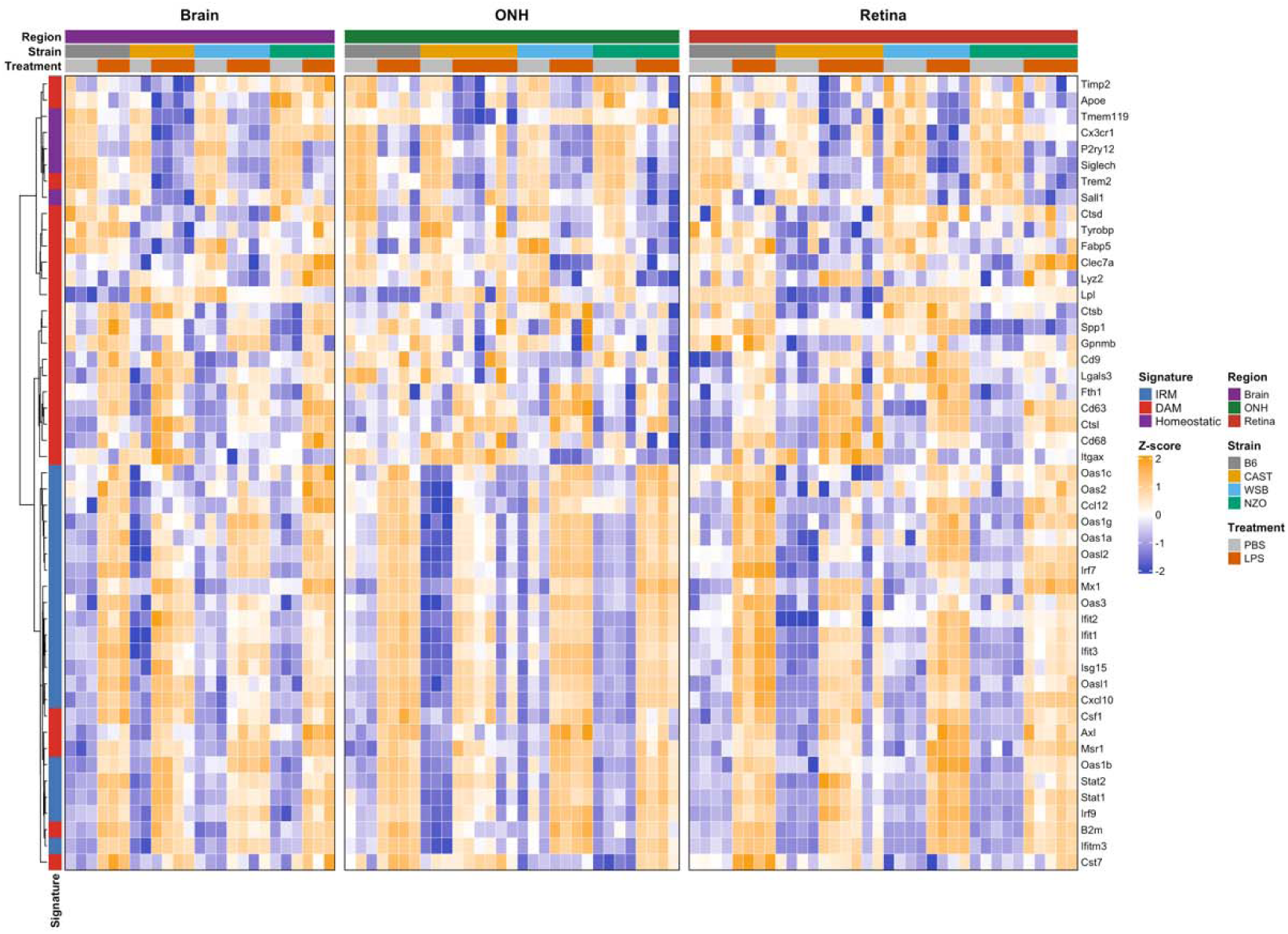
LPS challenge induces aspects of interferon-response microglia (IRM) and disease-associated microglia (DAM) gene expression across three CNS regions. Heatmap of 49 homeostatic, IRM, and DAM marker genes across brain, optic nerve head (ONH), and retina from four mouse strains (C57BL/6J, CAST/EiJ, NZO/HILtJ, WSB/EiJ) treated with LPS or PBS. Gene expression was quantified as TMM-normalized log-counts per million (log-CPM). Brain (B6: PBS n = 3, B6 LPS n = 3; CAST n= 2/4; NZO: n = 3/3; WSB n = 3/4). ONH (B6: n = 3/4; CAST: n = 3/6; NZO: n = 4/4; WSB: n = 3/4). Retina (B6: n = 4/4; CAST: n = 4/6; NZO: n = 5/5; WSB: n = 4/4)

Taken together, these data demonstrate that genetic background substantially modulates the transcriptional response to systemic LPS across CNS regions, with the retina showing the greatest strain-dependent transcriptional divergence from B6. The region- and strain-specific responses provides context on how genetics influence the neuroinflammatory transcriptional response to LPS.

To enable the broader research community to leverage these data, we have developed a web-based application for querying and visualizing differential expression and pathway enrichment results across all tissue, strain, and treatment comparisons [20]. As an example, we queried the expression of the putative murine ortholog of the glaucoma-risk lncRNA *CDKN2B-AS1*, *Gm12610*, in this dataset and found that LPS induced elevated expression of this lncRNA and was highest in ONH (Supplemental Figure 2). Altogether, this resource is intended to support hypothesis generation for studies investigating neuroinflammatory mechanisms in diseases of the brain and eye, the contribution of genetic context to inflammatory vulnerability, and the development of targeted therapeutic strategies.

## Conclusions

Neuroinflammation is a central driver of pathology in diseases of both the brain and the eye, yet how systemic inflammatory insults differentially affect distinct CNS compartments remains poorly understood. Systemic inflammation can disrupt the blood-brain and blood-retina barriers promoting neurodegenerative processes and is influenced by cellular environment and genetic context [21]. Bacterial products such as lipopolysaccharide (LPS) potently activate innate immune signaling and have been widely used to model systemic inflammation in preclinical studies [22]. However, the extent to which LPS-induced transcriptional responses are shared versus tissue-specific across CNS compartments, and how genetic background shapes these responses, has not been systematically examined.

Our transcriptomic profiling revealed that tissue identity is the dominant axis of variation within our dataset. This finding underscores that despite their shared CNS origin and proximity; these three compartments are molecularly distinct environments. Yet, LPS induced a conserved core of 1444 DEGs across all three tissues, enriched for innate immunity and acute-phase response pathways. This conserved signature demonstrates that systemic LPS elicits a shared neuroinflammatory response across the CNS, likely reflecting the disruption of the blood-brain and blood-retina barriers permitting peripheral inflammatory signals to reach neural tissue [23, 24].

Beyond the conserved core response, most LPS-induced changes were tissue-specific. The ONH showed the largest transcriptional response of the three tissues, with 9510 DEGs and a nearly even split between upregulated and downregulated genes. Genes uniquely upregulated in the ONH were enriched for proteostasis-related pathways, including proteasome and ribosome biogenesis, whereas uniquely downregulated genes were enriched for metabolic pathways such as fatty acid metabolism, lipid catabolism, and pyruvate metabolism. These pathways are relevant to ONH astrocyte responses described in glaucoma models. In experimental glaucoma, Ribotag-mediated astrocyte-specific RNA-seq further showed that ONH astrocytes upregulate proteolysis and oxidative phosphorylation, with relatively limited neuroinflammatory changes [25]. This is consistent with the proteostasis-dominant signature observed in the LPS-treated ONH. Whereas the downregulation of fatty acid and lipid metabolic pathways after LPS treatment also support claims of metabolic remodeling in ONH astrocytes during glaucoma. Mechanical stretch alters the astrocyte bioenergetics, increasing glycolytic activity and glutamine dependency [26]. Astrocyte networks redistribute metabolic resources in response to elevated IOP, which can leave the donating optic nerve more vulnerable to additional metabolic stress [17]. Reviews of glial metabolism in glaucoma similarly highlight disrupted lipid metabolism and fatty acid oxidation in astrocytes as potential contributors to retinal ganglion cell vulnerability [27, 28]. The overlap between LPS induced changes and glaucoma associated pathways, especially proteostasis activation and metabolic suppression, suggests that systemic inflammation may place a glaucoma-relevant stress on the ONH even without elevated intraocular pressure. This is relevant to glaucoma and other optic neuropathies, where ONH glial reactivity and axonal injury are central pathological features [29]. Future studies should test whether repeated or chronic systemic inflammatory insults lead to cumulative ONH damage and contribute to optic nerve disease.

A notable feature of our brain transcriptomic data is the relative paucity of pathway-level enrichment, both for the overall LPS response and for strain-differential responses, despite hundreds of differentially expressed genes. This contrasts with the retina and ONH, where coherent pathway-level signatures emerged. It is important to note that as we used whole hemibrain tissue for our transcriptomic analyses, the resulting signals average across structurally and cellularly heterogeneous regions, including the cortex, hippocampus, cerebellum, hypothalamus, and striatum. Multiple studies have demonstrated that systemic LPS elicits region-specific transcriptional responses in the brain, with distinct magnitudes and temporal dynamics across the hippocampus, hypothalamus, cortex, and cerebellum [30–32]. Furthermore, microglia the primary resident immune cells mediating the brain’s response to LPS exhibit region-dependent transcriptional identities and differential sensitivities to inflammatory stimuli [33, 34]. It is therefore possible that whole-tissue homogenization averages out region-specific programs that pull in opposing directions, diluting coherent pathway-level signals below the threshold for enrichment. This raises an alternate possibility that strain effects are broadly distributed across biological processes, such effects may instead be region-specific but hidden by tissue averaging. Future studies employing region-dissected or single-nucleus profiling of the brain following systemic LPS will be required to resolve these possibilities. Importantly, the present whole-brain data remain valuable for identifying the conserved, brain-wide component of the LPS response and for cross-tissue comparison with the retina and ONH, which are themselves regionally defined structures.

A central question in neuroinflammation is the extent to which genetic background shapes CNS vulnerability to inflammatory insults. We have previously demonstrated that genetic context modulates aging and degeneration in the murine retina across these same four strains [9]. Here, we extend this framework to the acute inflammatory setting and find that genetic background substantially modulates the transcriptional response to systemic LPS across all three CNS compartments, with the retina showing the greatest strain-dependent divergence. In the retina, WSB and NZO mice exhibited the largest strain-divergent responses relative to B6 (2115 and 1490 DEGs, respectively), while CAST mice showed a more modest divergence (639 DEGs). The retinal strain-divergent response in WSB mice was enriched for DNA replication and cell cycle pathways, while NZO mice showed enrichment for immune and metabolic processes. Given that WSB mice exhibit age-related photoreceptor loss and NZO mice, a model of metabolic syndrome, exhibit retinal ganglion cell loss and microvascular dysfunction [9], the strain-specific transcriptional responses to LPS identified here may reflect pre-existing differences in retinal cell type composition or cellular stress states that alter the inflammatory response. These findings raise the possibility that individuals with pre-existing retinal vulnerabilities may be disproportionately susceptible to inflammation-driven retinal injury supported by human epidemiological studies.

Population-based studies have linked chronic systemic inflammation with increased risk of ocular disease where surgery-indicated chronic rhinosinusitis a condition marked by persistent sinonasal inflammation, was associated with increased glaucoma risk [35], and HPV infection (Human Papillomavirus) was associated with a 34% higher hazard of incident glaucoma [36]. Elevated systemic inflammatory markers were also associated with greater glaucoma odds in the NHANES cohort (National Health and Nutrition Examination Survey) [37]. Similar associations extend beyond glaucoma: chronic hepatitis C infection was linked to increased risk of nonexudative AMD (Age-related Macular Degeneration) in a nationwide Taiwanese study [38], and systemic immune-mediated inflammatory diseases, including rheumatoid arthritis, systemic lupus erythematosus, and Crohn’s disease, have been associated with increased AMD risk [39]. Experimental work further shows that early-life infections can reprogram retinal microglia and worsen neovascular AMD later in life [40], while COVID-19 cohort studies have reported persistent retinal microvascular and ONH changes up to 12 months post-infection [41, 42]. Together, these findings suggest that systemic inflammation may heighten susceptibility to retinal injury, particularly in individuals with pre-existing ocular vulnerability.

The ONH showed the smallest strain-divergent response overall, with WSB exhibiting 415 DEGs, CAST 309 DEGs, and NZO only 106 DEGs relative to B6. Differentially responsive LPS DEGs in WSB mice in the ONH were enriched for extracellular matrix and structural remodeling pathways, consistent with the known role of ONH astrocytes in mediating structural responses to stress. The relatively constrained strain-divergent response in the ONH, despite its large overall LPS response, suggests that the core ONH inflammatory program is broadly conserved across genetic backgrounds, with genetic context modulating a smaller subset of tissue-remodeling and structural responses.

In totality, our efforts: 1) generated a high-depth transcriptomic dataset profiling the response to systemic inflammation across three CNS compartments in four genetically diverse mouse strains to facilitate hypothesis development and refinement, 2) identified a conserved innate immune core response shared across all three tissues, 3) determined that tissue identity is the dominant axis of variation with the ONH exhibiting proteostasis and metabolic pathway shifts and the retina showing suppression of phototransduction, 4) demonstrated that genetic background substantially modulates the inflammatory response, with the retina showing the greatest strain-dependent divergence, and 5) identified strain-specific transcriptional vulnerabilities that may reflect pre-existing differences in cellular stress states.

## Author contributions

MM, KDO, RLT, and GRH conceptualized the research. KDO and KJK performed LPS injections and tissue collection. TT and MM performed data analysis. MM and TT compiled and critically revised the first version. All authors read, edited, and approved the final version.

## Data Availability

Bulk RNA-seq have been deposited at GEO (Accession Pending) and are publicly available as of the date of publication. A shiny app for querying these data sets is available at: https://thejacksonlaboratory.shinyapps.io/Howell_LPS_Explorer/

## References

[1] H. Jung et al., “LPS induces microglial activation and GABAergic synaptic deficits in the hippocampus accompanied by prolonged cognitive impairment,” Sci Rep, vol. 13, no. 1, p. 6547, Apr 21 2023, doi: 10.1038/s41598-023-32798-9.

[2] S. Lehnardt et al., “Activation of innate immunity in the CNS triggers neurodegeneration through a Toll-like receptor 4-dependent pathway,” Proc Natl Acad Sci U S A, vol. 100, no. 14, pp. 8514–9, Jul 8 2003, doi: 10.1073/pnas.1432609100.

[3] H. A. Quigley, E. M. Addicks, W. R. Green, and A. E. Maumenee, “Optic nerve damage in human glaucoma. II. The site of injury and susceptibility to damage,” Arch Ophthalmol, vol. 99, no. 4, pp. 635–49, Apr 1981, doi: 10.1001/archopht.1981.03930010635009.

[4] H. A. Quigley and E. M. Addicks, “Chronic experimental glaucoma in primates. II. Effect of extended intraocular pressure elevation on optic nerve head and axonal transport,” (in eng), Invest Ophthalmol Vis Sci, vol. 19, no. 2, pp. 137–52, Feb 1980.

[5] M. A. Margeta et al., “Apolipoprotein E4 impairs the response of neurodegenerative retinal microglia and prevents neuronal loss in glaucoma,” Immunity, vol. 55, no. 9, pp. 1627–1644 e7, Sep 13 2022, doi: 10.1016/j.immuni.2022.07.014.

[6] E. H. Corder et al., “Gene Dose of Apolipoprotein E Type 4 Allele and the Risk of Alzheimer’s Disease in Late Onset Families,” Science, vol. 261, no. 5123, pp. 921–923, 1993, doi: doi:10.1126/science.8346443.

[7] E. J. Chesler et al., “The Collaborative Cross at Oak Ridge National Laboratory: developing a powerful resource for systems genetics,” Mamm Genome, vol. 19, no. 6, pp. 382–9, Jun 2008, doi: 10.1007/s00335-008-9135-8.

[8] G. A. Churchill, D. M. Gatti, S. C. Munger, and K. L. Svenson, “The Diversity Outbred mouse population,” Mamm Genome, vol. 23, no. 9-10, pp. 713–8, Oct 2012, doi: 10.1007/s00335-012-9414-2.

[9] O. J. Marola et al., “Genetic context modulates aging and degeneration in the murine retina,” Mol Neurodegener, vol. 20, no. 1, p. 8, Jan 20 2025, doi: 10.1186/s13024-025-00800-9.

[10] A. Dobin et al., “STAR: ultrafast universal RNA-seq aligner,” (in eng), Bioinformatics, vol. 29, no. 1, pp. 15–21, Jan 1 2013, doi: 10.1093/bioinformatics/bts635.

[11] J. Lilue et al., “Sixteen diverse laboratory mouse reference genomes define strain-specific haplotypes and novel functional loci,” Nat Genet, vol. 50, no. 11, pp. 1574–1583, Nov 2018, doi: 10.1038/s41588-018-0223-8.

[12] Y. Liao, G. K. Smyth, and W. Shi, “featureCounts: an efficient general purpose program for assigning sequence reads to genomic features,” Bioinformatics, vol. 30, no. 7, pp. 923–30, Apr 1 2014, doi: 10.1093/bioinformatics/btt656.

[13] Y. Liao, G. K. Smyth, and W. Shi, “The Subread aligner: fast, accurate and scalable read mapping by seed-and-vote,” (in eng), Nucleic Acids Res, vol. 41, no. 10, p. e108, May 1 2013, doi: 10.1093/nar/gkt214.

[14] P. Flicek et al., “Ensembl 2013,” Nucleic Acids Res, vol. 41, no. Database issue, pp. D48–55, Jan 2013, doi: 10.1093/nar/gks1236.

[15] M. D. Robinson, D. J. McCarthy, and G. K. Smyth, “edgeR: a Bioconductor package for differential expression analysis of digital gene expression data,” Bioinformatics, vol. 26, no. 1, pp. 139–40, Jan 1 2010, doi: 10.1093/bioinformatics/btp616.

[16] T. Wu et al., “clusterProfiler 4.0: A universal enrichment tool for interpreting omics data,” Innovation (Camb), vol. 2, no. 3, p. 100141, Aug 28 2021, doi: 10.1016/j.xinn.2021.100141.

[17] C. Gene Ontology et al., “The Gene Ontology knowledgebase in 2023,” (in eng), Genetics, vol. 224, no. 1, May 4 2023, doi: 10.1093/genetics/iyad031.

[18] A. Noailles, V. Maneu, L. Campello, P. Lax, and N. Cuenca, “Systemic inflammation induced by lipopolysaccharide aggravates inherited retinal dystrophy,” Cell Death Dis, vol. 9, no. 3, p. 350, Mar 2 2018, doi: 10.1038/s41419-018-0355-x.

[19] C. Collaborative Cross, “The genome architecture of the Collaborative Cross mouse genetic reference population,” Genetics, vol. 190, no. 2, pp. 389–401, Feb 2012, doi: 10.1534/genetics.111.132639.

[20] M. M. Tejas Temker, Kelly J. Keezer, Kristen D. Onos, Richard T. Libby, Gareth R. Howell. “Howell LPS Explorer.” https://thejacksonlaboratory.shinyapps.io/Howell_LPS_Explorer/ (accessed July 2, 2026.

[21] I. Galea, “The blood-brain barrier in systemic infection and inflammation,” Cell Mol Immunol, vol. 18, no. 11, pp. 2489–2501, Nov 2021, doi: 10.1038/s41423-021-00757-x.

[22] J. Sakai et al., “Lipopolysaccharide-induced NF-kappaB nuclear translocation is primarily dependent on MyD88, but TNFalpha expression requires TRIF and MyD88,” Sci Rep, vol. 7, no. 1, p. 1428, May 3 2017, doi: 10.1038/s41598-017-01600-y.

[23] D. Kokona, A. Ebneter, P. Escher, and M. S. Zinkernagel, “Colony-stimulating factor 1 receptor inhibition prevents disruption of the blood-retina barrier during chronic inflammation,” J Neuroinflammation, vol. 15, no. 1, p. 340, Dec 12 2018, doi: 10.1186/s12974-018-1373-4.

[24] C. Wei et al., “Brain endothelial GSDMD activation mediates inflammatory BBB breakdown,” Nature, vol. 629, no. 8013, pp. 893–900, 2024/05/01 2024, doi: 10.1038/s41586-024-07314-2.

[25] A. G. Mazumder, A. M. Jule, and D. Sun, “Astrocytes of the optic nerve exhibit a region-specific and temporally distinct response to elevated intraocular pressure,” Mol Neurodegener, vol. 18, no. 1, p. 68, Sep 27 2023, doi: 10.1186/s13024-023-00658-9.

[26] N. Pappenhagen, E. Yin, A. B. Morgan, C. C. Kiehlbauch, and D. M. Inman, “Stretch stress propels glutamine dependency and glycolysis in optic nerve head astrocytes,” (in English), *Front Neurosci*, Original Research vol. 16, p. 957034, 2022–August–05 2022, doi: 10.3389/fnins.2022.957034.

[27] A. Rombaut, R. Brautaset, P. A. Williams, and J. R. Tribble, “Glial metabolic alterations during glaucoma pathogenesis,” (in English), Frontiers in Ophthalmology, Review vol. Volume 3 - 2023, 2023–November–28 2023, doi: 10.3389/fopht.2023.1290465.

[28] Y. Hao, D. Liang, M. Ren, F. Kuang, and M. Wu, “Astrocyte Heterogeneity and Metabolic Reprogramming: Mechanisms Governing Retinal Ganglion Cell Damage in Glaucoma,” Cells, vol. 15, no. 6, p. 487, Mar 10 2026, doi: 10.3390/cells15060487.

[29] M. R. Hernandez, “The optic nerve head in glaucoma: role of astrocytes in tissue remodeling,” Prog Retin Eye Res, vol. 19, no. 3, pp. 297–321, May 2000, doi: 10.1016/s1350-9462(99)00017-8.

[30] C. Andre, J. C. O’Connor, K. W. Kelley, J. Lestage, R. Dantzer, and N. Castanon, “Spatio-temporal differences in the profile of murine brain expression of proinflammatory cytokines and indoleamine 2,3-dioxygenase in response to peripheral lipopolysaccharide administration,” J Neuroimmunol, vol. 200, no. 1-2, pp. 90–9, Aug 30 2008, doi: 10.1016/j.jneuroim.2008.06.011.

[31] H. A. Silverman et al., “Brain region-specific alterations in the gene expression of cytokines, immune cell markers and cholinergic system components during peripheral endotoxin-induced inflammation,” Mol Med, vol. 20, no. 1, pp. 601–11, Mar 11 2015, doi: 10.2119/molmed.2014.00147.

[32] H. Noh, J. Jeon, and H. Seo, “Systemic injection of LPS induces region-specific neuroinflammation and mitochondrial dysfunction in normal mouse brain,” Neurochem Int, vol. 69, pp. 35–40, Apr 2014, doi: 10.1016/j.neuint.2014.02.008.

[33] K. Grabert et al., “Microglial brain region-dependent diversity and selective regional sensitivities to aging,” Nat Neurosci, vol. 19, no. 3, pp. 504–16, Mar 2016, doi: 10.1038/nn.4222.

[34] E. Brandi et al., “Brain region-specific microglial and astrocytic activation in response to systemic lipopolysaccharides exposure,” (in English), Front Aging Neurosci, Original Research vol. 14, p. 910988, 2022–August–26 2022, doi: 10.3389/fnagi.2022.910988.

[35] S. F. Chau et al., “The Development of Glaucoma after Surgery-Indicated Chronic Rhinosinusitis: A Population-Based Cohort Study,” Int J Environ Res Public Health, vol. 16, no. 22, p. 4456, Nov 13 2019, doi: 10.3390/ijerph16224456.

[36] W. C. Liao et al., “Human Papillomavirus Infection Is Associated with Increased Risk of Glaucoma: Evidence in an Electronic Health Record Study,” Ophthalmology, vol. 132, no. 10, pp. 1161–1168, Oct 2025, doi: 10.1016/j.ophtha.2025.05.019.

[37] X. Li, Y. Q. Sun, X. D. Zhong, Z. J. Zhang, J. F. Tang, and Z. Y. Luo, “Association between systemic inflammatory response index and glaucoma incidence from 2005 to 2008,” (in English), Front Med (Lausanne), Original Research vol. 12, p. 1542073, 2025–February–04 2025, doi: 10.3389/fmed.2025.1542073.

[38] C. C. Yeh et al., “Increased Risk of Age-Related Macular Degeneration with Chronic Hepatitis C Virus Infection: A Nationwide Population-Based Propensity Score-Matched Cohort Study in Taiwan,” Viruses, vol. 13, no. 5, p. 790, Apr 28 2021, doi: 10.3390/v13050790.

[39] P. Shukla et al., “Propensity-Matched Analysis of the Risk of Age-Related Macular Degeneration with Systemic Immune-Mediated Inflammatory Disease,” Ophthalmol Retina, vol. 8, no. 8, pp. 778–785, Aug 2024, doi: 10.1016/j.oret.2024.01.026.

[40] M. Hata et al., “Early-life peripheral infections reprogram retinal microglia and aggravate neovascular age-related macular degeneration in later life,” (in eng), J Clin Invest, vol. 133, no. 4, Feb 15 2023, doi: 10.1172/JCI159757.

[41] K. Jevnikar et al., “The Comparison of Retinal Microvascular Findings in Acute COVID-19 and 1-Year after Hospital Discharge Assessed with Multimodal Imaging-A Prospective Longitudinal Cohort Study,” Int J Mol Sci, vol. 24, no. 4, p. 4032, Feb 17 2023, doi: 10.3390/ijms24044032.

[42] M. Ozturk, D. Kumova Guler, E. E. Oskan, and F. Onder, “Long-Term Effects of COVID-19 on Optic Disc and Retinal Microvasculature Assessed by Optical Coherence Tomography Angiography,” Diagnostics (Basel), vol. 15, no. 1, p. 114, Jan 6 2025, doi: 10.3390/diagnostics15010114.

