## Supplemental File 1 and 2 for "Genetic context alters central nervous system compartment dependent responses to lipopolysaccharide"

**Supplemental Figure**

**Figure 01**

**Transcriptomic profiling of LPS-induced systemic inflammation across three neural tissues.** MDS plots of RNA-seq samples colored by A tissue brain n=25, ONH n=31, retina n=36), B treatment (LPS vs. PBS; brain: LPS n=14, PBS n=11; ONH: LPS n=18, PBS n=13; retina: LPS n=19, PBS n=17), and C mouse strain (C57BL/6J, CAST/EiJ, NZO/HlLtJ, WSB/EiJ).

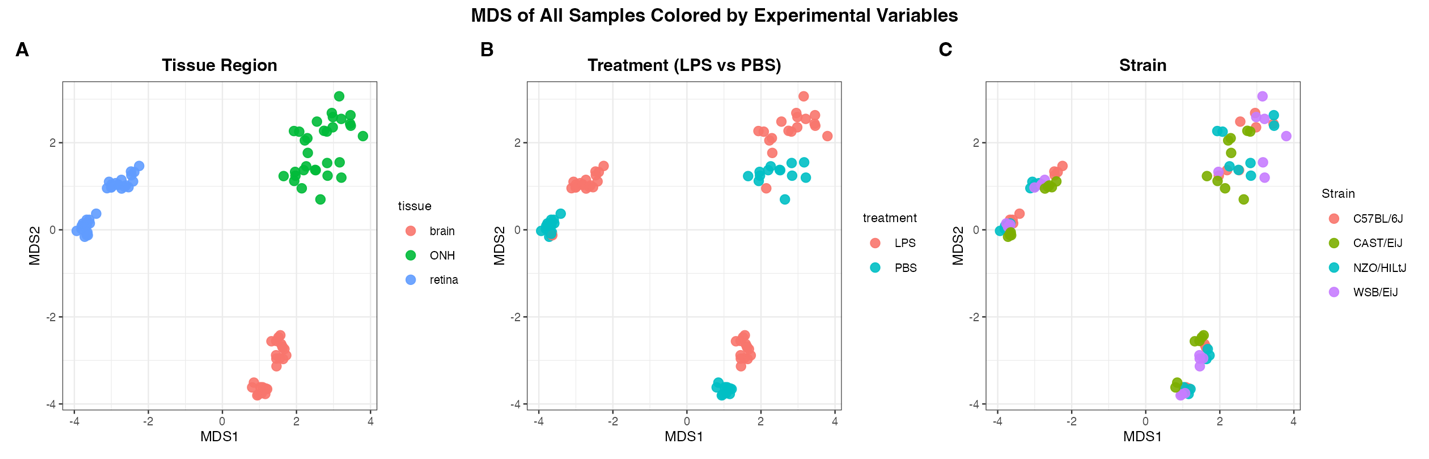

**Figure 02**

**LPS challenge induces *Gm12610* expression across three CNS regions.** Boxplot of *Gm12610* across brain, optic nerve head (ONH), and retina from C57BL/6J mice. Gene expression was quantified as TMM-normalized log-counts per million (log-CPM). Created using the web-app located at: “https://thejacksonlaboratory.shinyapps.io/Howell_LPS_Explorer/”. Brain (n = 3/group). ONH (B6: n = 3 PBS / 4 LPS). Retina (B6: n = 4/group)

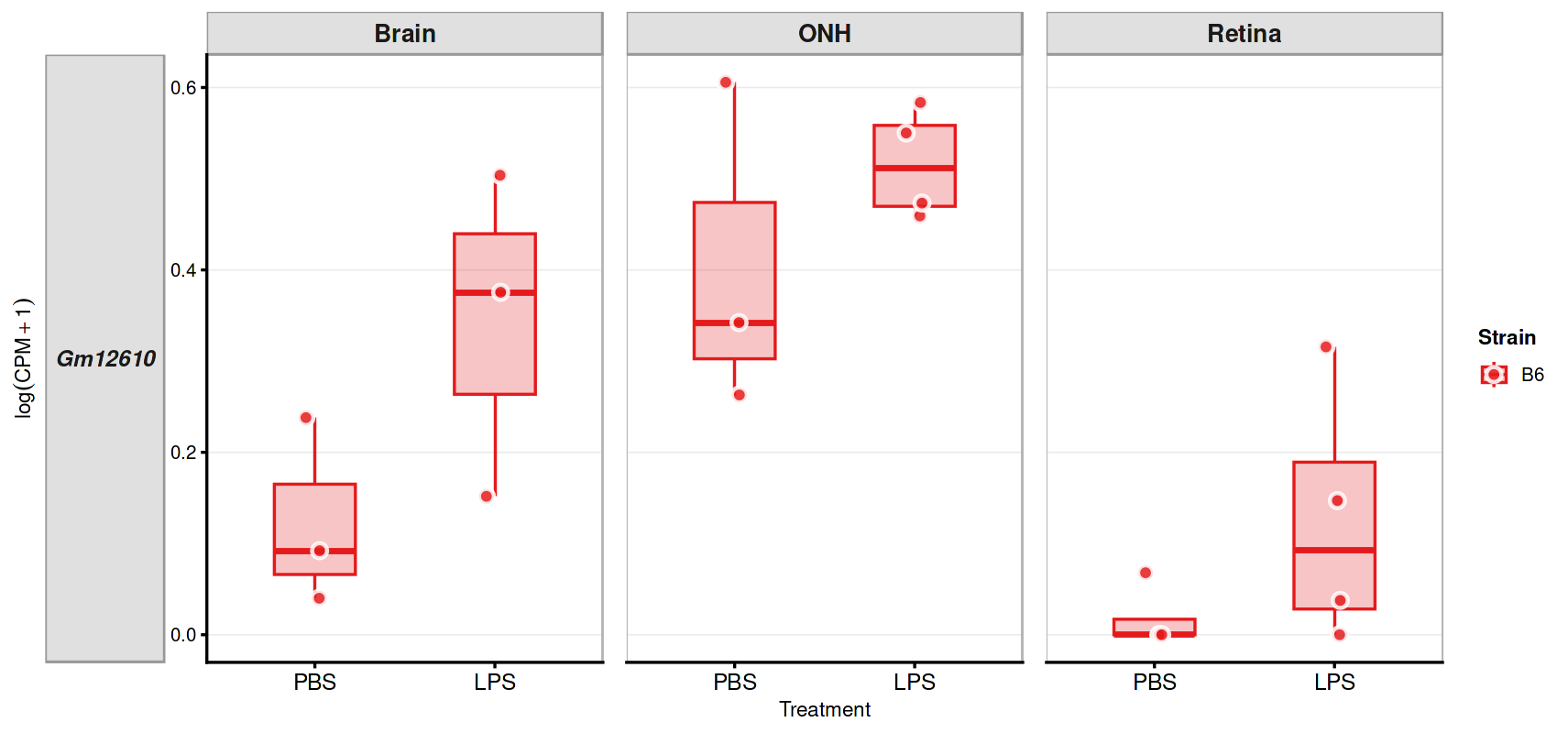
